# A predominately pulmonary activation of complement in a mouse model of severe COVID-19

**DOI:** 10.1101/2024.05.31.596892

**Authors:** Peter J. Szachowicz, Christine Wohlford-Lenane, Cobey J. Heinen, Shreya Ghimire, Biyun Xue, Timothy J. Boly, Abhishek Verma, Leila MašinoviĆ, Jennifer R. Bermick, Stanley Perlman, David K. Meyerholz, Alejandro A. Pezzulo, Yuzhou Zhang, Richard J.H. Smith, Paul B. McCray

## Abstract

Evidence from in vitro studies and observational human disease data suggest the complement system plays a significant role in SARS-CoV-2 pathogenesis, although how complement dysregulation develops in patients with severe COVID-19 is unknown. Here, using a mouse-adapted SARS-CoV-2 virus (SARS2-N501Y_MA30_) and a mouse model of severe COVID-19, we identify significant serologic and pulmonary complement activation following infection. We observed C3 activation in airway and alveolar epithelia, and in pulmonary vascular endothelia. Our evidence suggests that while the alternative pathway is the primary route of complement activation, components of both the alternative and classical pathways are produced locally by respiratory epithelial cells following infection, and increased in primary cultures of human airway epithelia in response to cytokine exposure. This locally generated complement response appears to precede and subsequently drive lung injury and inflammation. Results from this mouse model recapitulate findings in humans, which suggest sex-specific variance in complement activation, with predilection for increased C3 activity in males, a finding that may correlate with more severe disease. Our findings indicate that complement activation is a defining feature of severe COVID-19 in mice and lay the foundation for further investigation into the role of complement in COVID-19.

## Introduction

The COVID-19 pandemic has resulted in >7 million deaths worldwide (https://covid19.who.int/). Through a global scientific effort, we have learned much about SARS-CoV-2 and COVID-19 pathophysiology. COVID-19 exists on a spectrum from mild disease, characterized by minimal vague and generalized symptoms, to severe illness, with the most critically ill suffering from a combination of acute respiratory distress syndrome (ARDS) and multi-organ failure (1). Although the advent of effective vaccines and antiviral treatments has reduced hospitalizations and death (2–7), treatment for patients presenting with respiratory failure remains ineffective (5–7). With the ongoing threat of virulent mutations and pathogenic variants (8, 9), research that improves our understanding of this disease and identifies novel therapeutic options remains critical.

Reported pathophysiologic features of severe COVID-19 include prothrombotic complications (10–13), “cytokine storm” (14, 15), NETosis (16), and excessive cytotoxic T-cell responses (17), all which can be driven by complement activation. Complement is an ancient and evolutionarily conserved arm of the immune system (18). While its function in the innate immune response is well described, recent research suggests complement may play a role in cell-mediated and humoral adaptive immune responses (19). Additionally, several non-canonical functions of complement have been identified including roles in cell metabolism, autophagy, and apoptosis (20–22). While most commonly understood as liver-derived proteins, some cell types produce complement proteins locally (23, 24), with the lungs representing a large reservoir of extrahepatic complement production (25). In the context of infections, complement is a key mediator of the host response to respiratory viruses, including other pathologic human coronaviruses (26–28). In particular, SARS-CoV-2 is associated with complement dysregulation locally and systemically. Genomic and proteomic studies have confirmed autopsy results demonstrating COVID-19 patients have an increased complement response compared to ARDS from other causes, and patients with underlying disorders characterized by complement overactivation are prone to developing more severe disease than matched cohorts (29–32). Thus, evidence supports the idea that SARS-CoV-2 infection may result in complement activation and this may contribute to disease severity.

Despite evidence for complement involvement in COVID-19, significant gaps remain in our understanding of its contributions to this disease. For example, there is no consensus on which complement pathway is activated, with studies implicating the classical (33), lectin (34), or alternative pathways (35–37), and whether SARS-CoV-2 antigens activate complement directly or indirectly by stimulating inflammatory and immune mediators remains uncertain (34, 36, 37). Respiratory epithelial cells do generate complement in response to cell injury and inflammation (38, 39) and this response has been demonstrated in cultured primary human alveolar type II cells following SARS-CoV-2 infection (37). However, confirmatory human and animal model data are lacking, and therefore, the impact of complement on COVID-19 disease pathogenesis remains undefined.

To address this knowledge gap, we investigated the role of complement in a mouse model of COVID-19 that recapitulates many features of severe disease in humans. We demonstrate that following infection, complement activation is detected in both serum and lungs as early as 2 days post-infection (dpi). This complement activation occurs primarily via the alternative pathway and is specific to the lungs, with mice developing an extensive complement burden and C3 colocalization in alveoli, airways, and blood vessels. Importantly, we also show complement deposition is not strictly limited to infected respiratory epithelial cells, suggesting activation of complement extends beyond direct antiviral mechanisms. This study provides the first demonstration in an animal model of severe COVID-19 that complement protein C3 is locally produced and activated in the lung where it may play a direct role in disease pathogenesis.

## Results

### SARS2-N501Y_MA30_ reproduces the clinical phenotype of severe COVID-19

We previously reported the development of a mouse model of COVID-19 that recapitulates features of severe, human disease (40). To investigate the complement response, we infected young (6-8 week old) BALB/c mice intranasally with 5,000 PFU of SARS2-N501Y_MA30_ virus, and euthanized mice on 0-5 dpi for sample collection. A second group was monitored for weight loss and mortality. All virus-treated mice succumbed to infection by 7 dpi (Figure 1A) and lost ∼30% of their body weight (Figure 1B). Interestingly, we observed a sex specific response to infection and noted a trend towards improved survival for females (Supplementary figure 1), consistent with other murine models of SARS-CoV-2 infection (41). In evaluating respiratory viral burden, we saw peak SARS2-N501Y_MA30_ N protein (nucleocapsid protein) antigen staining by 2 dpi, with evidence of viral clearance by 4 dpi (Figures 1C and D); a pattern confirmed with viral titer by plaque assay (Figure 1E). In all, the mice met metrics indicative of severe disease.

**Figure 1.**
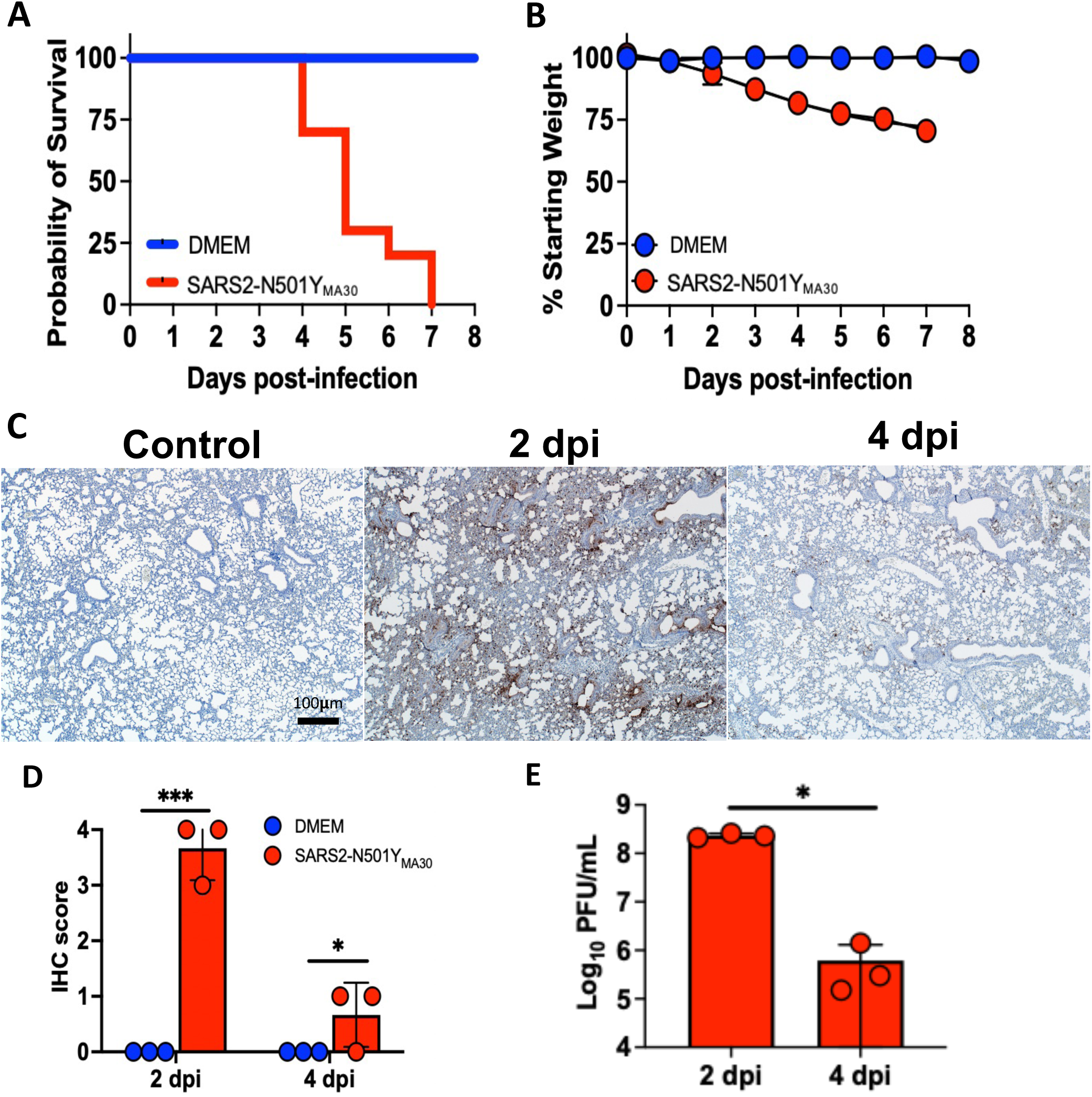
BALB/c mice infected with SARS2-N501Y_MA30_ develop severe disease with peak respiratory viral burden 2 dpi. *(A)* Probability of survival following a 5,000 PFU inoculum of SARS2-N501Y_MA30_ (red) compared to control group (blue) with significance achieved by 5 dpi, p<0.05 (n=10 mice per group). *(B)* Mean +/- SEM weight loss post-infection assessed by percent decrease from initial weight at each time point with significant difference achieved by 3 dpi, p<0.05 (n=10 mice per group). *(C)* Immunohistochemical (IHC) staining for SARS2-N501Y_MA30_ nucleocapsid protein (N-protein) (brown = N protein, n=3 mice per group). *(D)* IHC scoring for severity by percent area of lung stained positive for viral antigen and *(E)* viral titer obtained from plaque assay on lung homogenates (n=3 mice per group). Red = SARS2-N501Y_MA30_ and blue = DMEM (significance determined by log-rank test and unpaired t-test, where * indicates p<0.05 and *** indicates p<0.0001). DMEM = Dulbecco’s Modified Eagle’s Medium.

### Complement response following infection with SARS2-N501Y_MA30._

We hypothesized that, like humans with COVID-19, mice infected with SARS2-N501Y_MA30_ would mount a complement response in the lungs. To test this hypothesis, lung homogenates were collected from infected mice at 2 and 4 dpi, which correspond with the peak viral burden (Figures 1C-E) and peak illness severity before death, respectively. To evaluate complement cascade activity (Figure 2), we quantified factor B (FB) (alternative pathway specific), C3 (common cascade) and C4 (classical and lectin pathway specific) proteins in lung tissue (Figure 3). Pulmonary C3 and C3b were elevated at 4 dpi on both western blot (Figure 3A) and ELISA (Figure 3D and E), suggesting C3 convertase is formed in lung tissue and highly active following infection. Alternative pathway specific FB and Bb were also significantly elevated on 4 dpi (Figure 3B). While we detected C4 (Figure 3C), it was not significantly increased compared to controls. Data for 2 dpi demonstrate a similar pattern suggesting C3 and FB activation post-infection, with no significant change in lung C4 levels (Supplementary figure 2). Of note, we did not observe consistent differences between male and female mice on C3 or C4 western blot, though the C3b ELISA showed a trend toward increased levels in males on 2 and 4 dpi.

**Figure 2.**
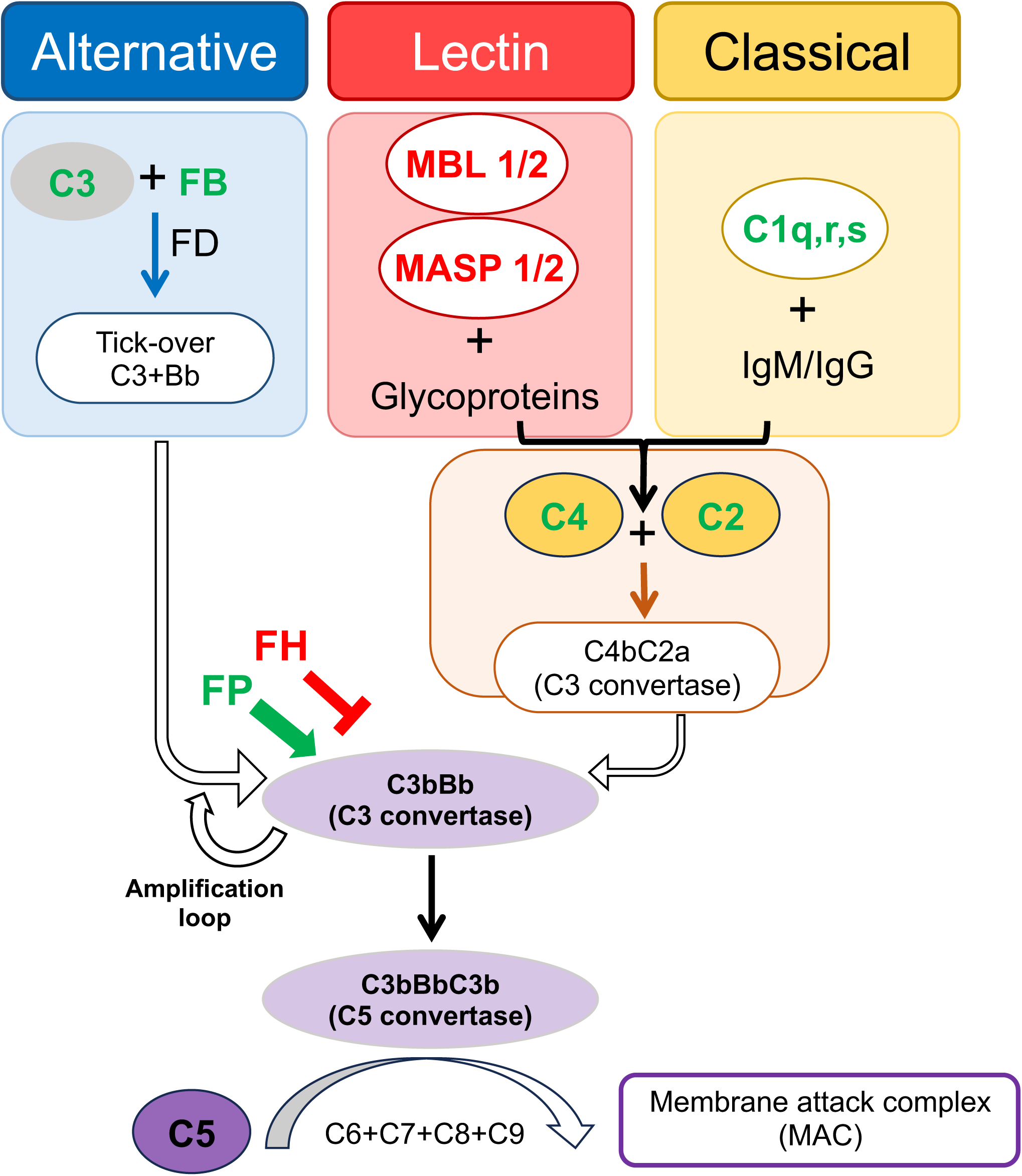
Schematic of complement cascade and components upregulated by SARS-CoV-2 infection. Green text indicates proteins with increased mRNA transcripts in mice following SARS2-N501Y_MA30_ infection, red text indicates proteins with reduced mRNA transcript abundance in mice following SARS2-N501Y_MA30_ infection. As depicted, the alternative pathway is continuously and spontaneously activated at low levels (tick-over) to generate C3bBb (C3 convertase), though rapid self-enhancement occurs by feedback amplification loop once activation is initiated. The lectin and classical pathways have different triggering substrates (glycoproteins and immunoglobulins, respectively), but both converge on the generation of C4 and C2, which join to create the C3 convertase C4bC2a. Although the lectin and classical pathways are initiated via separate mechanisms, a majority of their activity ultimately converges on C3bBb (C3 convertase), which cleaves C3 into C3a (anaphylatoxin) and C3b (opsonin). All three pathways coalesce with C5 convertase cleavage of C5, and the generation of C5a (anaphylatoxin) and C5b (joins with C6-C9 to form C5b-9, membrane attack complex). In addition, there are various cofactor proteins that help regulate cascade activity, such as properdin (FP) and complement factor H (FH), which can augment or inhibit further activation, respectively. FB = Factor B. MBL = mannose binding lectin. MASP = MBL-associated serine protease. FD = Factor D.

**Figure 3.**
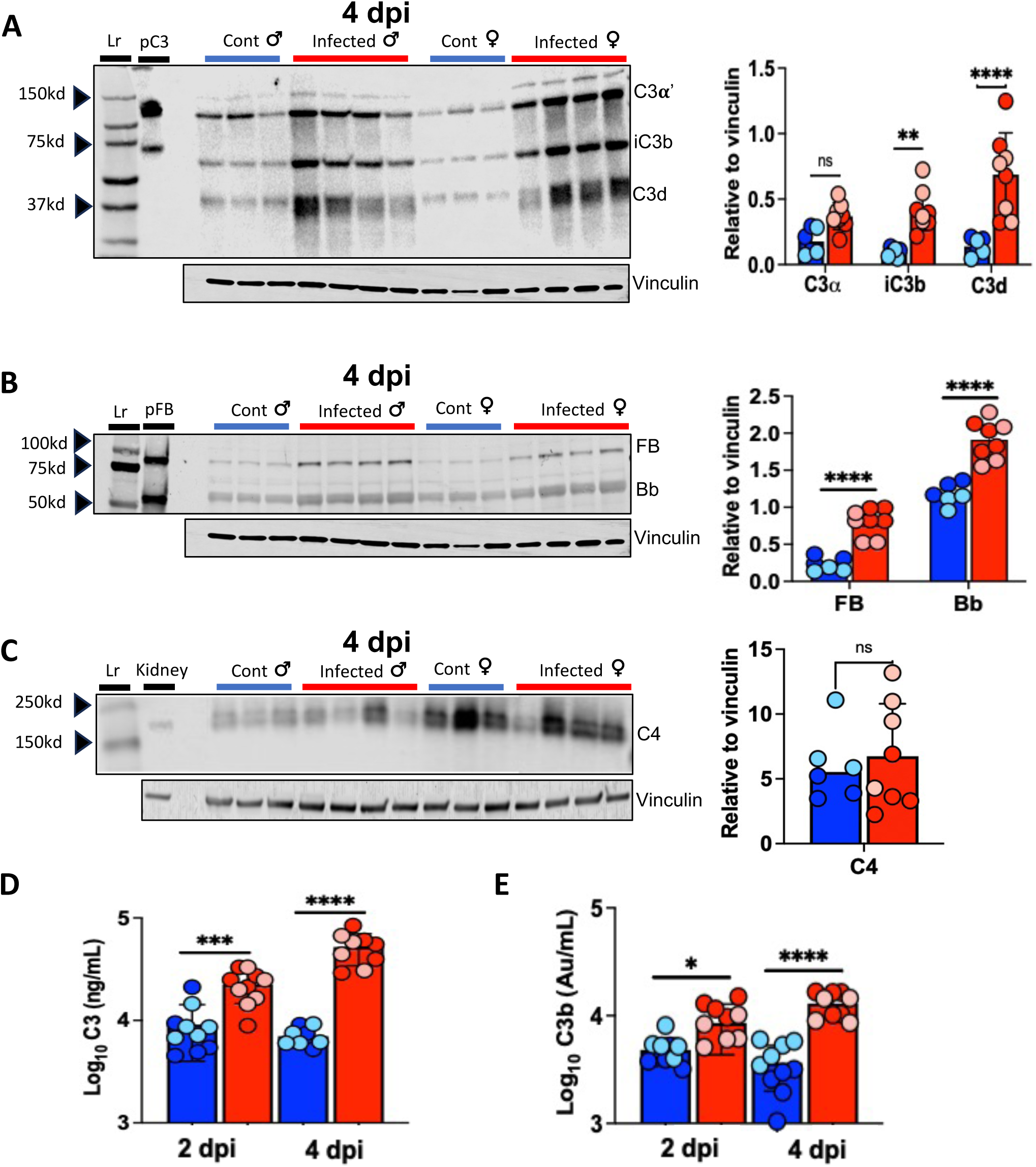
Increased C3 activation in lungs of mice infected with SARS2-N501Y_MA30_. Western blot of lung homogenates with corresponding densitometry (right) demonstrates increases in iC3b and C3d *(A)* and FB and Bb (*B*), but not C4 *(C)* on 4 dpi. Purified human C3 (pC3), purified Factor B and Bb (pFB), and uninfected mouse kidney homogenates were used as controls for C3, FB, and C4, respectively. *(D* and *E)* Lung homogenate ELISA for C3 and C3b demonstrates increases with SARS2-N501Y_MA30_ infection compared to control lung at 2 and 4 dpi. Blue bars indicate samples from control mice, red bars indicate samples from infected mice, light blue and light red circles indicate females, dark blue and dark red circles indicate males. n = 10 mice per group. Significance as determined by one-way ANOVA and unpaired t-test, * p<0.05,** p<0.001, *** p<0.0001, **** p<0.00001. Lr = ladder.

We hypothesized that C3 would co-localize to cells infected with SARS2-N501Y_MA30_. To test this hypothesis, we immunolocalized the viral N protein and C3 in infected tissues and observed that while C3 co-localized with N protein at 2 and 4 dpi, C3 also deposited in cells with no detectable N protein (Figure 4A), suggesting potential indirect mechanism of C3 activation.

**Figure 4.**
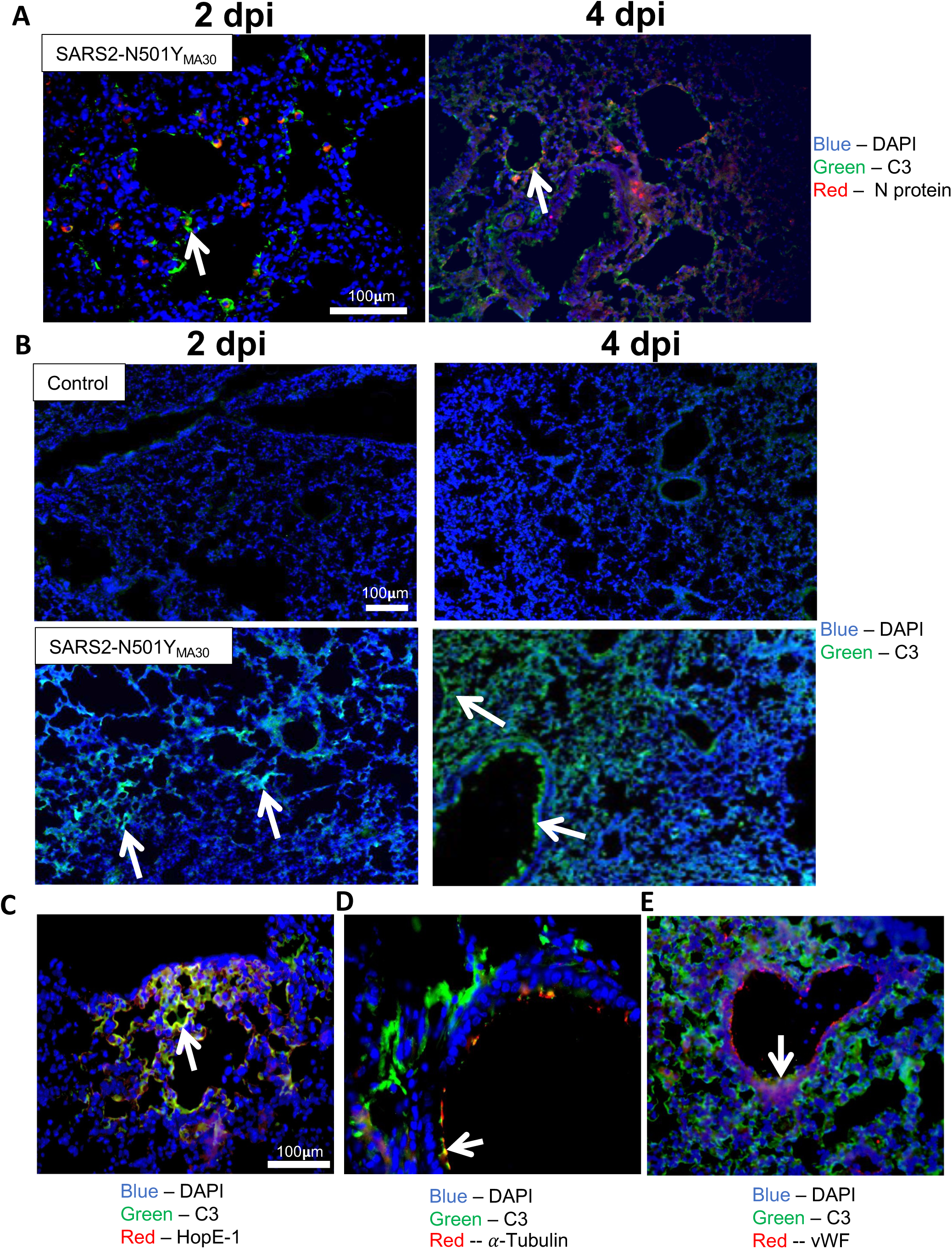
Complement C3 is colocalized with cells in various pulmonary sub compartments. (*A)* Colocalization of C3 (green) and SARS2-N501Y_MA30_ nucleocapsid protein (red) on 2 and 4 dpi. n= 5, blue = DAPI, red = nucleocapsid protein, green = C3, white arrows indicate areas with C3 and nucleocapsid protein colocalization. *(B)* Staining for complement C3 on lung tissue from control (top row) and following infection with SARS2-N501Y_MA30_ (bottom row) on 2 and 4 dpi. White arrows indicate areas of C3 staining. n = 5, green = C3, blue = DAPI. *(C)* Colocalization of C3 with alveolar type I cells, *(D)* ciliated airway cells, and *(E)* endothelial cells on 4 dpi. White arrows indicate regions of colocalization with C3. dpi = days post-infection, vWF = von Willebrand Factor.

To identify the cell types responsible for pulmonary C3 activation, we immunolocalized C3 in lung tissue sections. C3 expression was evident by 2 dpi, with notable C3 deposition as early as 1 dpi (Supplementary figure 3), with extensive involvement by 4 dpi (Figure 4B). C3 expression was diffuse, with localization to airways, alveoli, and blood vessels. Of note, these regions of the lung are common sites of injury in COVID-19 patients with small airways disease, diffuse alveolar damage, and pulmonary vascular thrombosis (42, 43).

Notably, in lung tissue C3 localized to cells at lumenal surfaces (Figure 4B and C). By co-staining for C3 and HopE-1 (type I alveolar cell marker), acetylated α-tubulin (ciliated cell marker), and von Willebrand factor (vWF, endothelial cell marker), we found C3 co-localized with these cell types along their apical membranes, confirming C3 associates with cell types in multiple sub-compartments in the lungs, including those where little or no viral infection occurred (Figure 4A). The apical polarization of C3 also suggests the presence of membrane-bound convertase formation on the lumenal cell surface. Of note, we also localized C4 in the lungs of infected mice, but expression did not change substantially over the course of infection, consistent with immunoblot data.

These results are consistent with extensive C3 activation in the lung following SARS2-N501Y_MA30_ infection, and suggest C3 convertase formation by activation by the alternative pathway in the absence of any significant change in classical or lectin pathway activity.

### Local complement production by respiratory epithelial cells

Although the observed pulmonary C3 could be explained by deposition of circulating complement, we suspected an alternate etiology. Extrahepatic complement production by respiratory epithelial cells has been described (44) and given the abundance of complement detected in lung tissue, we hypothesized respiratory epithelial cells were sources of local production. To that end, we performed RNAscope for C3 and found RNA transcripts in respiratory epithelial cells in mice infected with SARS2-N501Y_MA30_ (Figure 5A). C3 and SARS2-N501Y_MA30_ RNA transcripts colocalized at 2 dpi (Figure 5A), confirming that infected cells express C3. However, C3 mRNA was also observed in SARS2-N501Y_MA30_-negative cells consistent with the above findings that uninfected cells are also sources for C3 in the lungs. We hypothesized these uninfected cells were generating complement in response to cytokines released by neighboring cells. Respiratory epithelial cells produce IL-17 and TNF-α in response to various respiratory viral infections including SARS-CoV-2 (45–49), and these cytokines, among others (i.e., IL-1β and IL-6), can induce complement synthesis (50, 51). This suggested the hypothesis that a paracrine or systemic cytokine response may trigger respiratory epithelial cell C3 activation. Thus, we exposed primary cultures of HAE to a cocktail of IL-17 and TNF-α and demonstrated the upregulation of C3 and numerous genes involved in the complement cascade, further supporting this as a mechanism which might explain our findings (Supplementary figure 4).

**Figure 5.**
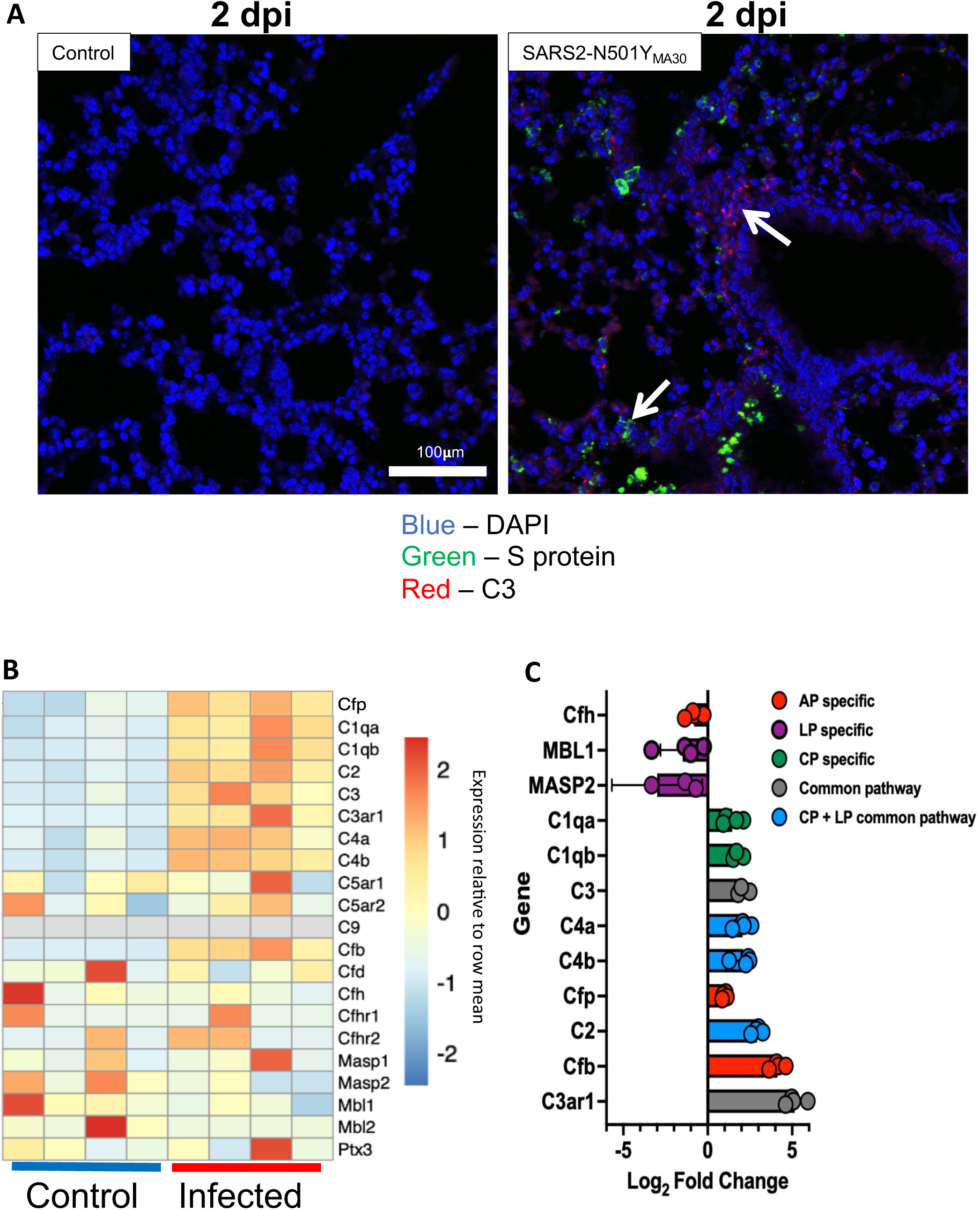
Messenger RNA expression analysis suggests local production of complement proteins in lung tissue following SARS2-N501Y_MA30_ infection. *(A)* RNAscope for S protein RNA and C3 RNA from lung section on 2 dpi following 5,000 PFU of SARS2-N501Y_MA30_. White arrows indicate C3 RNA and SARS2-N501Y_MA30_ S protein RNA, red = C3, green = S protein, and blue = DAPI. *(B)* Heat map depicting the results of bulk RNA sequencing for genes associated with complement pathways on lung tissue collected 5 dpi follow inoculation with 1,000 PFU SARS2-N501Y_MA30_. *(C)* Mean log_2_ fold change in transcript abundance post-infection. Complement pathway indicated by bar color: red = alternative pathway specific genes, green = classical pathway specific genes, purple = lectin pathway specific genes, blue = shared lectin and classical pathway genes, grey = common pathway genes. n = 4. dpi = days post-infection.

To evaluate whether mice infected with SARS2-N501Y_MA30_ also had evidence increased transcription of complement genes other than C3, we performed bulk RNA sequencing of infected mouse lung tissue at 5 dpi. This showed increased expression of complement cascade mRNA transcripts compared with control animals (Figure 5B). Based on fold change, we observed significant increases in expression of *Cfb* (complement factor B) and *Cfp* (complement factor properdin) post-infection, both specific to the alternative pathway (Figure 5C). Contrary to protein analyses, both classical and alternative pathway RNA transcripts were increased. We observed decreased expression of *Masp1, Masp2, Mbl1,* and *Mbl2,* which are lectin pathway specific genes, as well as a decrease in *Cfh* (complement factor H) transcripts, a major regulator of complement activity. Of note, there was a large increase in *C3ar1* gene expression. C3aR is a receptor for the anaphylatoxin C3a, and its stimulation can result in the release of cytokines from epithelial and immune cells.

In summation, these findings suggest infection stimulates a widespread complement response that is driven by locally derived complement proteins. Both overactivation and impaired regulation of the alternative pathway contribute to a dysregulated complement response in animals following SARS2-N501Y_MA30_ infection.

### Systemic complement response following SARS2-N501Y_MA30_ infection

To evaluate whether systemic complement activity increases post-infection, we collected serum at 2 and 4 dpi and measured complement proteins by ELISA. Serum C3 levels in infected mice were increased compared to controls (Figure 6A), as was serum C4 (Figure 6B). CH_50_, used to assess classical and terminal complement pathway activity, was increased at 2 dpi but fell to control levels at 4 dpi (Figure 6C). Immunoblot for C3 protein on kidney homogenate of infected mice (Supplementary figure 5) showed no increase in C3 deposition, indicating the observed C3 response was limited to the lung. These findings suggest the major complement response is driven by local production of complement proteins.

**Figure 6.**
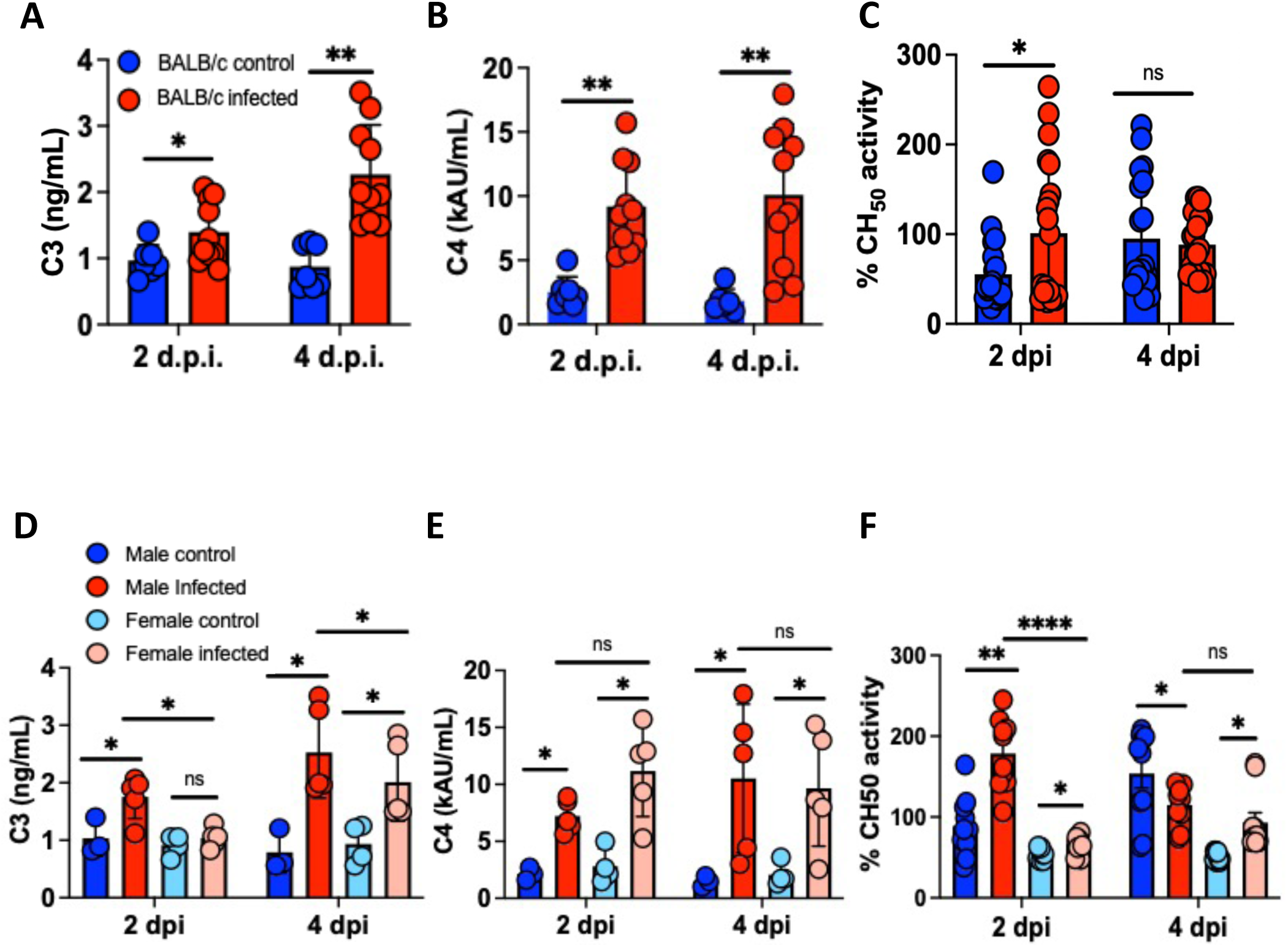
Characterization of systemic complement activity in mice infected with SARS2-N501Y_MA30_. *(A and B)* Serum C3 and C4 ELISA from control (blue) and infected (red) mice on 2 and 4 dpi. *(C)* Serum CH_50_ activity assay from control (blue) and SARS2-N501Y_MA30_ mice (red). Comparison of sex difference by serum ELISA for C3 *(D)* and C4 *(E)* on 2 and 4 dpi, with data from sex specific CH_50_ activity assay shown in *(F).* dpi = days post-infection. Significance determined by one-way ANOVA or unpaired t-test, * p<0.05, ** p<0.001, **** p<0.00001. ns = nonsignificant. C3 and C4 measured from n = 7. DMEM and n = 10 SARS2-N501Y_MA30_ mice per group. CH_50_ assay from n = 10 mice per group, samples run in duplicate.

As stated above, male mice had a more severe disease phenotype, a finding also described in human COVID-19 patients (52), prompting us to question whether there were sex differences in systemic complement responses to SARS2-N501Y_MA30_. Infected male mice had significantly increased serum C3 and CH_50_ activity compared to females (Figures 6D and 6F). Importantly, C4 expression post-infection was not significantly different between sexes (Figure 6E), indicating the observed difference in sex-specific complement activity is primarily via increased alternative pathway complement activation in response to SARS2-N501Y_MA30_ infection which correlated to trends observed in the lungs.

### Complement response contributes to lung pathology and inflammatory response

When evaluating the pattern of lung injury (Figure 7A), we noted that mice develop lung injury starting on 2 dpi, which further advanced by 4 dpi, well after the development of a significant complement response, but prior to substantial cytokine or immune cell infiltration (40). Complement components C3a and C5a are potent anaphylatoxins (53). To investigate the cytokine-chemokine profile in the setting of complement activation, we quantified concentrations of chemokines and cytokines in serum and lung homogenates at 0 to 5 dpi (Figure 7B). RANTES (CCL5), IL-1β, IL-6, CXCL-1, MCP-1, and CXCL-10 were elevated with infection, cytokines which have been associated with C3aR and C5aR1 stimulation (54–57). We assessed mouse lung inflammatory gene expression from the bulk RNAseq data at 5 dpi (Supplementary figure 6) and found increased transcription of genes associated with interferon signaling, as well as those typical of NLRP3 inflammasome and TLR mediated inflammation.

**Figure 7.**
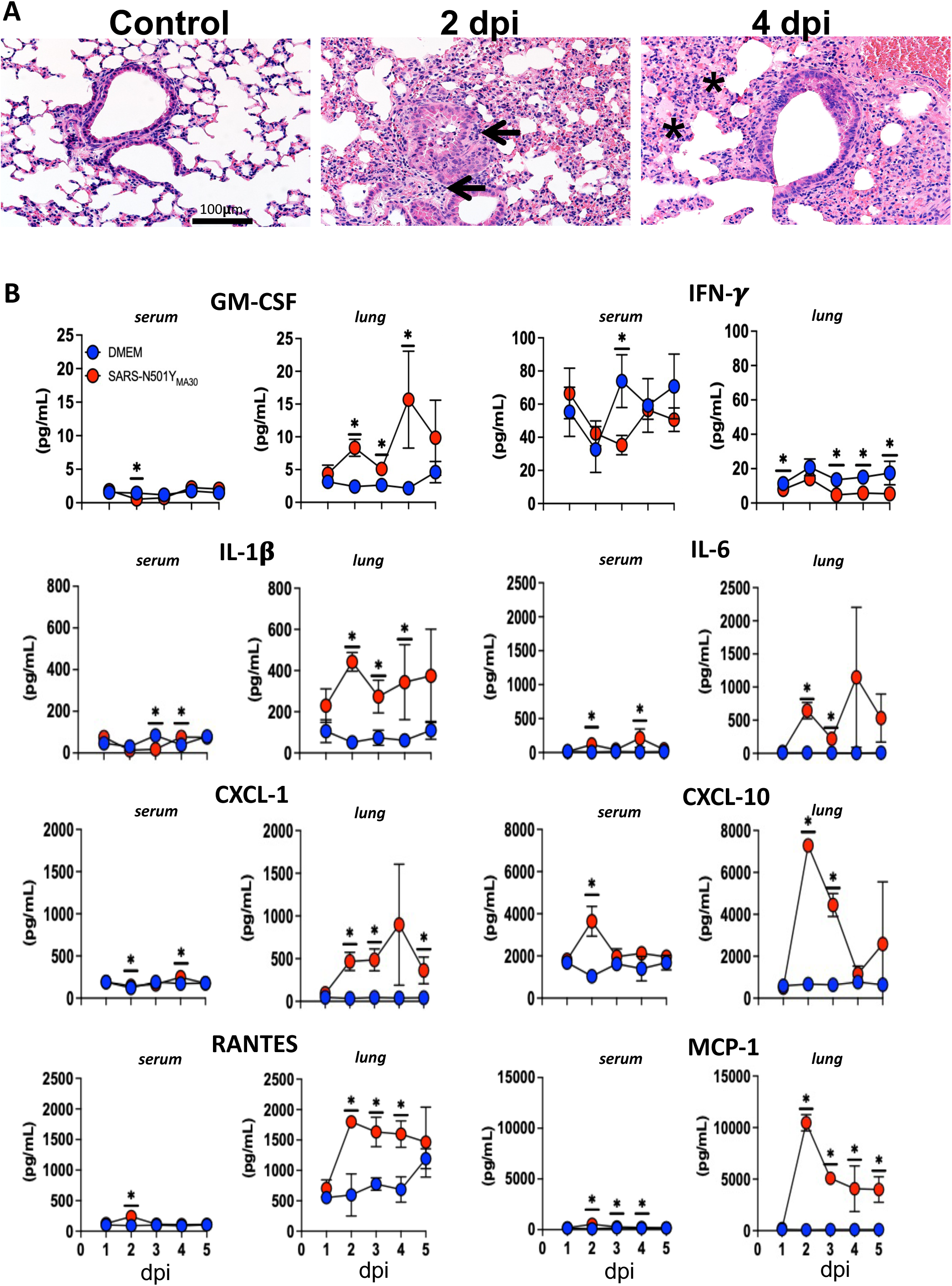
Lung histopathology and inflammatory response is consistent with complement mediated disease. *(A)* Representative images of H&E staining of lung slides from histopathologic exam, with control and infected mice on 2 and 4 dpi which demonstrates cellular infiltrates (2 dpi) (black arrows) and edema (4 dpi) (black asterisks). n = 3 mice per time point. *(B)* Serum and lung cytokine profiling for 1-5 dpi, title above graph in each column indicates whether sample is from serum or lung homogenate. n = 3 mice per group at each time point. DMEM (Dulbecco’s Modified Eagle’s Medium), dpi = days post-infection. Significance determined by unpaired t-test, * p < 0.05.

## Discussion

This study provides the first in-depth description of complement activity in an animal model of severe COVID-19. Our observations offer insights into an important aspect of this disease, with potential implications for other infectious and non-infectious forms of acute lung injury. Key findings include: 1) in response to SARS2-N501Y_MA30_ infection, complement activity is significantly increased in the lung (locally) and circulation (systemically); 2) this activity is in part locally derived from respiratory epithelial cells; 3) complement activation occurs primarily through the alternative pathway; 4) complement production and activation by respiratory epithelial cells is not limited to virus infected cells; 5) complement contributes to the observed COVID-19 pathophysiology by causing lung injury and inflammation.

Early in the course of infection, we observed significant C3 deposition in the lung, which becomes more diffuse by 4 dpi. The C3 is apically polarized and involves lumenal membranes of cells in the airways, alveoli, and blood vessels. These cells form the subcompartments of the lung that are primary areas of injury in COVID-19 ARDS. The detection of significant C3b levels shows that C3 convertase is formed and the complement cascade is active. Infected mice also had evidence of a systemic increase in complement activity based on C3 and C4 ELISA and CH_50_ activity assays, though we found no evidence of substantial complement activation in organs outside the lungs. Overall, these findings suggest that SARS-CoV-2 infection primarily induces a lung-specific complement response.

Whereas our studies, as well as others, demonstrate local production of complement by respiratory epithelial cells (37, 39, 44), no in vivo model of SARS-CoV-2 has confirmed this finding. Here, we provide evidence that mice infected with SARS2-N501Y_MA30_ have increased transcription of complement-activating genes and decreased expression of complement-regulating genes in the lung. *C3*, *Cfb, Cfp,* and *Cfh* are among the most significantly altered, suggesting alternative pathway activation in this model of COVID-19 (Figures 2 and 5). Importantly, this transcriptional data is supported by protein analysis as we observed pulmonary FB and Bb increases significantly post-infection (Figure 3B). We also observed increases in transcript abundance for genes associated with the classical pathway. However, a similar pattern of classical pathway upregulation was previously described in COVID-19 patients (35) and might reflect the later timepoint of 5 dpi for sample collection. This may coincide with the onset of immunocomplex formation and resultant IgM- and IgG-mediated activation of the classical pathway with subsequent C4 cleavage and C3 convertase formation (58). Thus, classical pathway activation may be a later event rather than the primary driver of complement activation and respiratory disease pathology. Likewise, the increase in serum C4 could reflect its role in the inflammatory response, with liver-derived C4 produced as a non-specific acute phase reactant rather than a specific response to SARS2-N501Y_MA30_ infection. Additionally, we note that while there was no overall significant increase in pulmonary C4, some male mice appeared to have higher C4 levels compared to controls on 2 dpi, and taken with the RNA sequencing data, future studies should explore this discrepancy. Together, these findings support the concept of a locally derived complement response, which may drive lung injury in COVID-19 and warrants future investigation. Additionally, identification of the triggering pathway has important mechanistic and therapeutic implications, providing grounds for future research.

Previous studies have suggested SARS-CoV-2 activates complement through direct interaction between viral antigen and complement proteins, or as a cellular response to viral invasion (36, 37). However, our results suggest that complement activation may be induced in a more paracrine or systemic fashion. The significance of this observation is that the complement response is diffuse, involves the entire lung, and induces widespread pulmonary inflammation and cellular injury. Consistent with this pathophysiology, the cytokine profile is one of inflammation which can be driven by the potent anaphylatoxins C3a and C5a binding to their respective receptors as described above (59–61). The pattern of gene expression could be consistent with a complement-mediated response via mitochondrial antiviral-signaling protein (MAVS) (57) in which NF-KB and Interferon regulatory factor 3 (IRF3) can be stimulated by C3 (57). A similar cytokine/chemokine pattern can also be associated with C3a and C5a stimulation of direct (C3aR1 and C5aR1) and indirect (potentiation of TLR signal) p38 MAPK inflammatory signalling, inducing NF-KB and activator protein 1 (AP-1) mediated cytokine/chemokine response (61). This suggests that following infection with SARS2-N501Y_MA30_, there is complement activation with a subsequent complement-mediated inflammatory response. However, these findings are hypothesis generating and further studies are required to understand the links between complement activation and the inflammatory response. This inflammation is primarily pulmonary in origin, consistent with our observation that lung tissue had the greatest complement activation following infection.

Humans and mice exhibit sex-specific differences in serologic complement responses (62); whether this baseline difference in alternative pathway activity contributes to the more severe disease phenotype in males deserves more investigation. Overall, these findings support a role of locally produced complement driving both lung injury and inflammation in COVID-19.

Our study has limitations. First, while this murine model recapitulates several features of COVID-19, mice are imperfect surrogates for humans and differ significantly in features of pulmonary anatomy, immune system function, and complement activity (63–65). Second, the data presented come from a single mouse strain, BALB/c. We know that mouse strains respond differently to viral infections, including SARS-CoV-2 (40), and the complement response can have inter-strain variability (66). Third, our experiments focused on a severe disease phenotype and followed mice over a short period as all infected animals succumbed to infection prior to day 10. Thus, we cannot extrapolate our findings to more mild iterations of disease. Likewise, we are unable to comment on the effect of complement activation in the recovery period or any implications this activation might have on the development and perpetuation of “long-COVID” (67). Finally, because our studies focused on describing the mechanism and role of complement activation in a mouse model of COVID-19, we did not explore the impact a deficit of complement, either by pharmaceutical intervention or genetic alteration, has on disease.

In summary, our findings provide strong evidence for local, respiratory epithelial cell derived complement activation in a mouse model of COVID-19. This activation occurs primarily via the alternative pathway. It is neither dependent on direct interaction with viral antigen, nor is it simply a cellular response to viral invasion. Our observation that C3 activation precedes any cytokine or cellular immune response in the lung supports the concept that both the local and systemic inflammatory response in COVID-19 are mediated by local complement activation. This work will serve as a foundation for studies addressing the role of complement inhibitors in the treatment of COVID-19. The approach presented here could also be used to explore the role of complement in other forms of infectious and non-infectious acute and chronic lung injury.

## Methods

### Mice, human tissue, cells, and virus

#### Sex as a biological variable

This study examined male and female mice, and sex-dimorphic effects are reported. Our study examined cell cultures taken from male and female donors, and similar findings are reported for both sexes.

#### Ethics Statement

Primary airway epithelia from human donors were isolated from discarded tissue, autopsy, or surgical specimens. Cells were provided by The University of Iowa *In Vitro* Models and Cell Culture Core Repository. Information that could be used to identify a subject was not provided. All studies involving human subjects received University of Iowa Institutional Review Board approval (Protocol #230167). Mouse experimental protocols were reviewed and approved by the University of Iowa Institutional Animal Care and Use Committee (IACUC), in accordance with the National Institutes of Health guidelines.

#### Animal studies

Animal studies were approved by the University of Iowa Animal Care and Use Committee. Mice were maintained in the University of Iowa Animal Care Unit under standard dark/light cycles, with controlled temperature and humidity. BALB/c mice were obtained (Charles River Laboratories) and used at 6-8 weeks of age. Mice were randomly assigned to groups, with numbers sufficient to obtain statistical significance.

#### Human airway epithelial cells and cell line cultures

The University of Iowa *In Vitro* Models and Cell Culture Core cultured and maintained HAE as previously described (68). Briefly, following enzymatic disassociation of trachea and bronchus epithelia, the cells were seeded onto collagen-coated, polycarbonate Transwell inserts (0.4 μm pore size; surface area = 0.33 cm^2^; Corning Costar, Cambridge, MA). HAE were submerged in Ultroser G (USG) medium for 24 hours (37°C and 5% CO_2_) at which point the apical media is removed to allow cells to differentiate at an air-liquid interface. VeroE6 cells (ATCC CRL-1586) were grown in Dulbecco’s Modified Eagle’s Medium (DMEM) with 10% FBS. The SARS2-N501Y_MA30_ was generated as previously described (40) and propagated in VeroE6 cells with input matching confirmed by genetic sequencing.

#### SARS-CoV-2 Infection

Mice were anesthetized with ketamine/xylazine (87.5mg/kg and 12.5mg/kg) and inoculated intranasally with the indicated dosage of SARS2-N501Y_MA30_ in 50µl DMEM or DMEM alone. Mice were examined and weighed daily, and euthanized per institutional protocol if weight loss was >30%. At various time points, blood was collected from submandibular vein, mice were euthanized, and indicated organs were collected. SARS-CoV-2 work was conducted in a Biosafety level 3 (BSL3) laboratory.

#### Quantitative Histopathology and Immunohistochemistry

Lung tissues were fixed in 10% neutral buffered formalin (5-7 days), dehydrated through a progressive series of alcohol and xylene baths, paraffin-embedded into blocks, sectioned (∼4 µm) and stained with hematoxylin and eosin (H&E) stain. Tissues were evaluated by boarded veterinary pathologist and evaluated in a post-examination method of masking the pathologist to group assignment (69). SARS-CoV-2 immunohistochemistry and scoring was performed as previously described (40) with minor modification in SARS-CoV-2 primary antibody (rabbit polyclonal anti-N-protein, Sino Bio 40143-T62, 1:4000 x 15 minutes). Briefly, edema and immunohistochemistry ordinal scores were based on distribution of each parameter in lung sections 0 – none, 1 - <25%, 2 - <50%, 3 <75% and 4 - >75%.

#### Antibodies and immunolocalization

Immunostaining was performed as described previously (70). The following antibodies were used: SARS-CoV-2 N-protein (1:500, cat#40588-T62, Sino Biological), goat anti-mouse C3 antibody (1:1000, cat#55463, MP Biomedicals), donkey anti-goat Alexa Fluor 488 (1:600, cat#A32814, Invitrogen), anti-alpha-tubulin (1:200, cat#5335S, Cell Signaling), anti-HopE1 (1:100, cat#sc-398703, Santa Cruz), anti-vWF (1:500, cat.#AB7356, Millipore), and Alexa Fluor 546 phalloidin (1:600, cat#A22283, Invitrogen). Slides were mounted with Vectashield-DAPI (cat#H-1200, Vector Laboratories) and visualized with a Keyence BZ-X810 fluorescence microscope (Keyence Corporation of America).

#### Bioplex cytokine profiling

The levels of cytokines and chemokines in mouse serum and lung homogenates were determined using the Bio-Plex Multiplex Immunoassay System as described previously (Bioplex 200 machine, Biorad, Hercules, CA, USA) (71). Repeat freeze-thaw cycles were avoided to minimize protein degradation. The following cytokines and chemokines were measured using a Bioplex Express Kit according to the manufacturer’s directions (Biorad, Hercules, CA, USA): GM-CSF, IFNγ, IL-6, IL-1β, RANTES, MCP-1, CXCL-1, CXCL-10.

#### Virus Titers

Lungs were dissociated in 1X PBS using a Bead Mill homogenizer (Fisherbrand). Supernatants were serially diluted and added to VeroE6 cells, after 1 hour incubation, Incocula were replaced with 0.6% agarose containing 2%FBS in DMEM and after 4 days cells were fixed with 20% formaldehyde and stained with 1% crystal violet for plaque assay, with final titer quantified as PFU/mL (40)

#### Western Blot Analysis

Mouse organ homogenates were combined with 1% NP40 and protease inhibitors per BSL3 virus sterilization protocol and frozen. 20 μg protein was loaded onto gels and run under reducing (C3) or non-reducing (C4) conditions. Following electrophoresis and transfer to PVDF membranes, samples were blocked with 5% milk and incubated with primary antibody anti-mouse C3 (1:500, cat#55463, MP Biomedicals) or anti-mouse C4 (1:200, cat#ab11863, Abcam) overnight. The membrane was washed with Invitrogen Novex tricine SDS running buffer and incubated with HRP-conjugated secondary for 1 hour at room temperature, then developed using Supersignal West Pico Plus chemiluminescent kit (cat#34580, ThermoScientific). Protein signal was read using Odyssey Imager. Membranes were stripped using Restore Western Stripping Buffer (cat#21059, ThermoScientific) and process repeated with anti-vinculin antibody (1:10,000, cat#700062, Invitrogen) as loading control. Densitometry analysis was performed using Image-J.

#### Cytokine stimulation of human airway epithelial cells

Cultures of human airway epithelia (HAE) cells were propagated, plated, and matured as described above. Mature HAE cultures were treated with basolateral TNF-α (10ng/mL) and IL-17 (20ng/mL) daily and cells were harvested at 5 hours and 48 hours post-cytokine stimulation. Cells were snap frozen and cell lysates were combined with Trizol and immediately stored at -80°C for preservation, with single cell RNA sequencing performed as described below.

#### Mouse lung bulk RNA sequencing

RNA was extracted and purified from mouse lungs using the RNeasy plus mini-kit (QIAGEN). A cDNA library was prepared using the Illumina TruSeq stranded mRNA protocol. Library concentration and fragment size was measured using Qubit-HS and High-Sensitivity DNA Assay. The pooled library was sequenced on a Illumina NovaSeq 6000 SP Flowcell instrument for paired end reads. Library preparation, quality control, and sequencing were done by University of Iowa Genomics Facility.

Base calls were demultiplexed and converted to FASTQ format by the University of Iowa Genomics Facility. Fastq files were pseudo-aligned to the mouse reference genome (72). Mapped raw reads were reported in transcripts per million by Kallisto. Downstream analysis was summarized as gene-level estimates using tximport v1.10.1 in Rv3.5 (73). Differential gene expression was performed using iDEP v0.92 (74). RNA-Seq data are available at GSE249304.

#### Human airway epithelial cell RNA Sequencing

Bulk RNA-sequencing was performed in collaboration with the University of Iowa Genomics Division using the manufacturer’s recommended protocols. Briefly, 500 ng DNase I–treated total RNA was enriched for polyA-containing transcripts using beads coated with oligo(dT) primers. The enriched RNA pool was fragmented, converted to cDNA, and ligated to sequencing adaptors using the TruSeq stranded mRNA sample preparation kit (RS-122-2101, Illumina). The molar concentrations of the indexed libraries were measured using the 2100 Bioanalyzer (Agilent) and combined equally into pools for sequencing. The concentrations of the pools were measured with the Illumina Library Quantification Kit (KAPA Biosystems) and sequenced on the Illumina HiSeq 4000 genome sequencer using 75 bp paired-end SBS chemistry. Pseudoalignment of raw sequencing reads and quantification of transcript-level expression were obtained using Kallisto version 0.45.0 and human transcriptome reference GRCh38. p12 (67). Gene counts were imported into R, and differential expression tests were performed using DESeq2 version 1.22.2 (68). Further, gene expression modeling in DESeq2 accounted for the experimental design, acknowledging and correcting for paired control and treated samples for each donor. Changes in complement related gene products were visualized as heatmaps generated using the Clustvis tool (https://biit.cs.ut.ee/clustvis/) (69). RNA-Seq data are available in the NCBI’s GEO database (GEO GSE 176121).

#### RNAscope

RNAscope was performed using the Advanced Cell Diagnostics protocol (ACD, Neward, CA). Fixed frozen paraffin-embedded lung sections were deparaffinized in xylene and 100% ethanol, then dried. Target retrieval was performed using RNAscope Target Retrieval Reagents (ACD, cat#322000), following H_2_O_2_ then protease treatment using RNAscope H_2_O_2_ and Protease Plus (ACD, cat#322330). For in-situ hybridization slides were incubated with RNA probes for SARS-CoV-2 S (RNAscope Probe V-nCoV-2-S-C2) and C3 (RNAscope Probe Mm-C3-C3), then mounted with DAPI Mounting Media (ACD).

#### Statistics

Statistics were performed using Graph Pad Prism 10 Software. Tests included unpaired or paired 2-tailed Student’s *t* test for comparing 2 groups, 1-way ANOVA with multiple comparison test for comparing more than 2 groups. Tests used are described in figure legend. P-value of <0.05 was considered significant.

#### Study Approval

All human and animal studies were approved by the University of Iowa Instituional Review Board.

## Data Availability

The Supporting Data Values XLS file includes values for each data point presented in the paper. GEO accession numbers for databases are included above. Further data supporting the findings of this study are available from the corresponding authors upon reasonable request.

## Author Contributions

PJS, CWL, PBM, RJHS, and YZ conceived and designed studies. PJS, AV, CJH, SG, AP, LM, and CWL performed experiments. CJH performed ELISA experiments. SG, BX, and AP performed RNA sequencing and data analysis. JRB and TJB contributed to cytokine assay. DKM performed histopathology and viral IHC. AV performed RNAscope. SP provided valuable insights in discussions and helped design experiments. PJS, PBM, YZ, and RJHS analyzed data. PJS, RJHS, and PBM wrote the manuscript. All authors approved of the manuscript.

## Acknowledgements

We would like to thank Paul Morgan, Wioleta Zelek, Ron Taylor, Santiago Rodriguez de Cordoba, and Matthew Pickering for the sharing of reagents, experimental assistance, and helpful discussions that contributed to this manuscript. We would like to thank the Tully family for their support. This work was supported by the National Institutes of Health (NIH) [P01 AI-060699], the University of Iowa Carver College of Medicine, and the Tully Family Foundation. PBM is supported by the Roy J. Carver Charitable Trust. Carver College of Medicine COVID-19 Grant (PBM and AP), and NHLBI 1R01HL163024 (AP).

Address correspondence to Peter J. Szachowicz, Department of Internal Medicine, 169 Newton Road, 6312A PBDB, Iowa City, Iowa 55242, USA. Phone: 319-384-1107. or Paul B. McCray, Jr., 169 Newton Road, 6320 PBDB, Iowa City, Iowa 52242, USA. Phone: 319.335.6844.

