## Supplemental figures 1-6 for "A predominately pulmonary activation of complement in a mouse model of severe COVID-19"

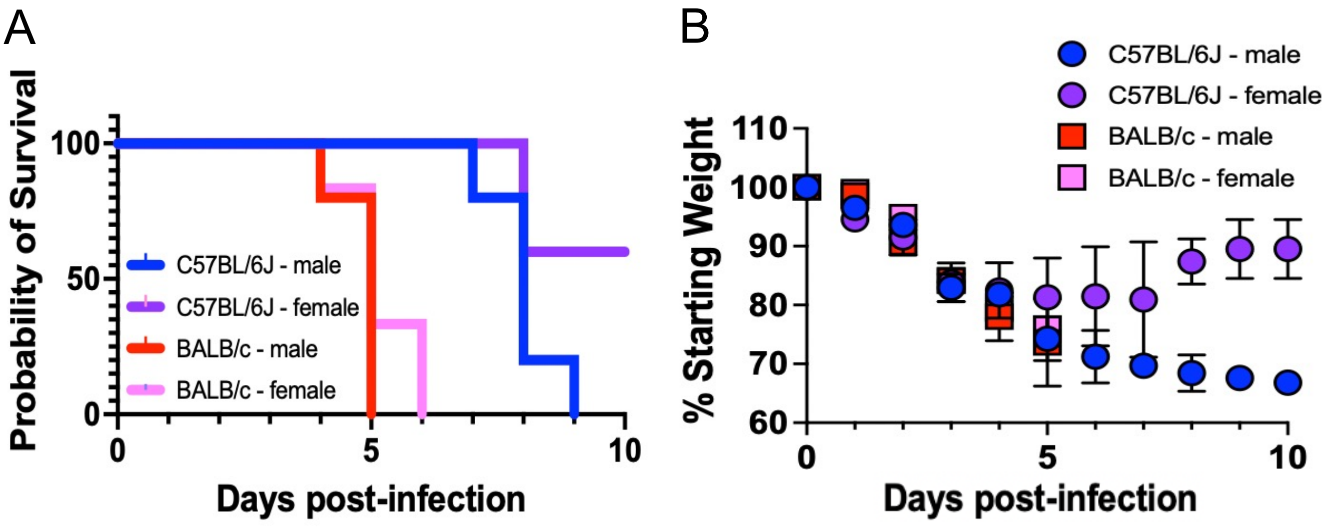


**Supplementary figure 1.** Sex differences in weight loss and mortality following SARS2-N501Y$_{MA30}$ infection is impacted by mouse strain. *(A)* Percent weight loss for male and female BALB/c and C57BL/6J mice show significant differences between male and female C57BL/6J mice by 8 dpi ($p<0.005$), but not in BALB/c mice. *(B)* Percent survival shows significant advantage for C57BL/6J mice compared to BALB/c mice of both sexes ($p<0.05$) and female C57BL/6J mice compared to male C57BL/6J mice ($p<0.05$), with a trend towards survival advantage in female BALB/c mice compared to male BALB/c mice ($p<0.308$). n = 6 mice per group. Statistical significance determined by unpaired t-test and log-rank test.

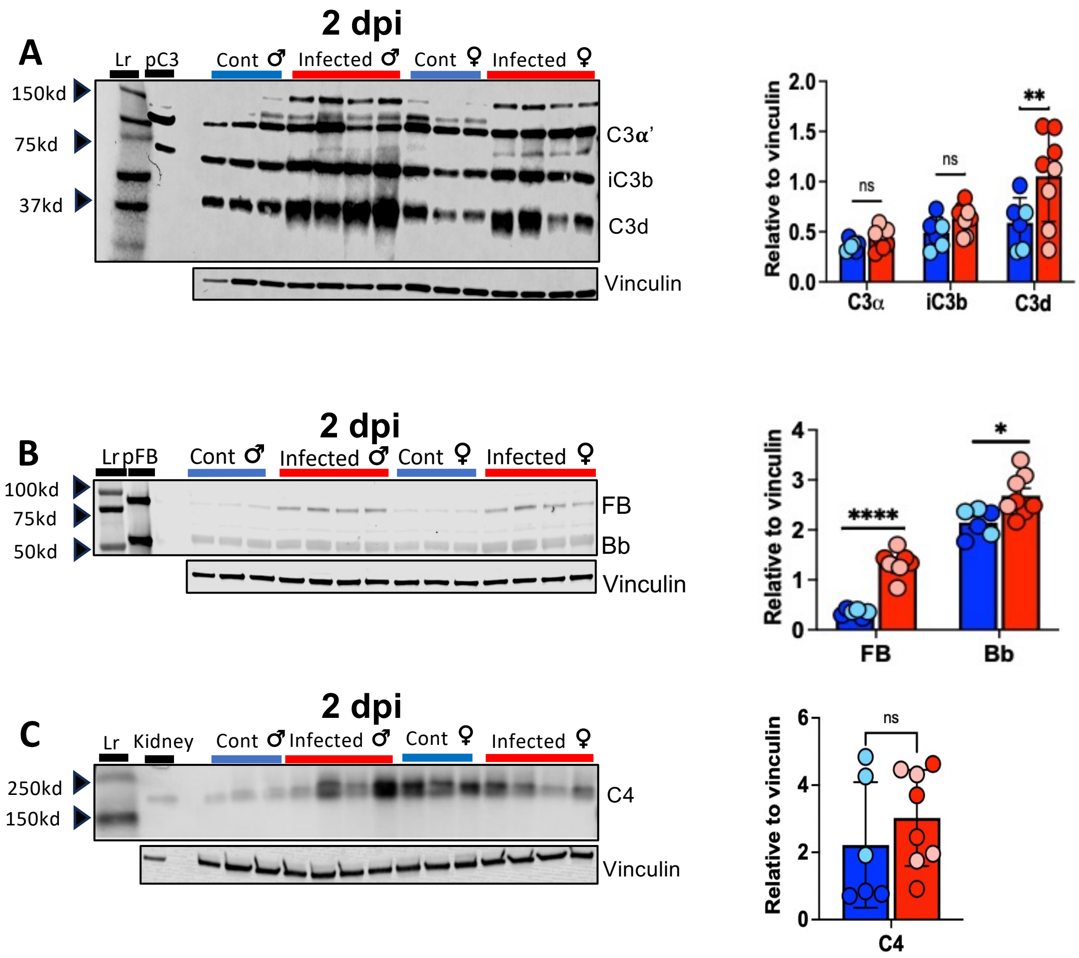


**Supplementary figure 2.** Complement FB and C3 are increased in lungs 2 days post-infection. Increased C3 activation in lungs of mice infected with SARS2-N501Y$_{MA30}$. Western blot of lung homogenates with corresponding densitometry (right) demonstrates increases in C3b and C3d *(A)* and FB and Bb *(B)* but not C4 *(C)* on 2 dpi. Purified human C3 (pC3), purified FB and Bb (pFB), and control mouse kidney homogenates were used as controls for C3, FB, and C4, respectively. Blue bars indicate samples from control mice, red bars indicate samples from infected mice, light blue and light red circles indicate females, dark blue and dark red circles indicate males. n = 10 mice per group. Significance as determined by one-way ANOVA, where * p<0.05, ** p<0.001, *** p<0.0001, **** p<0.00001. Lr = ladder.

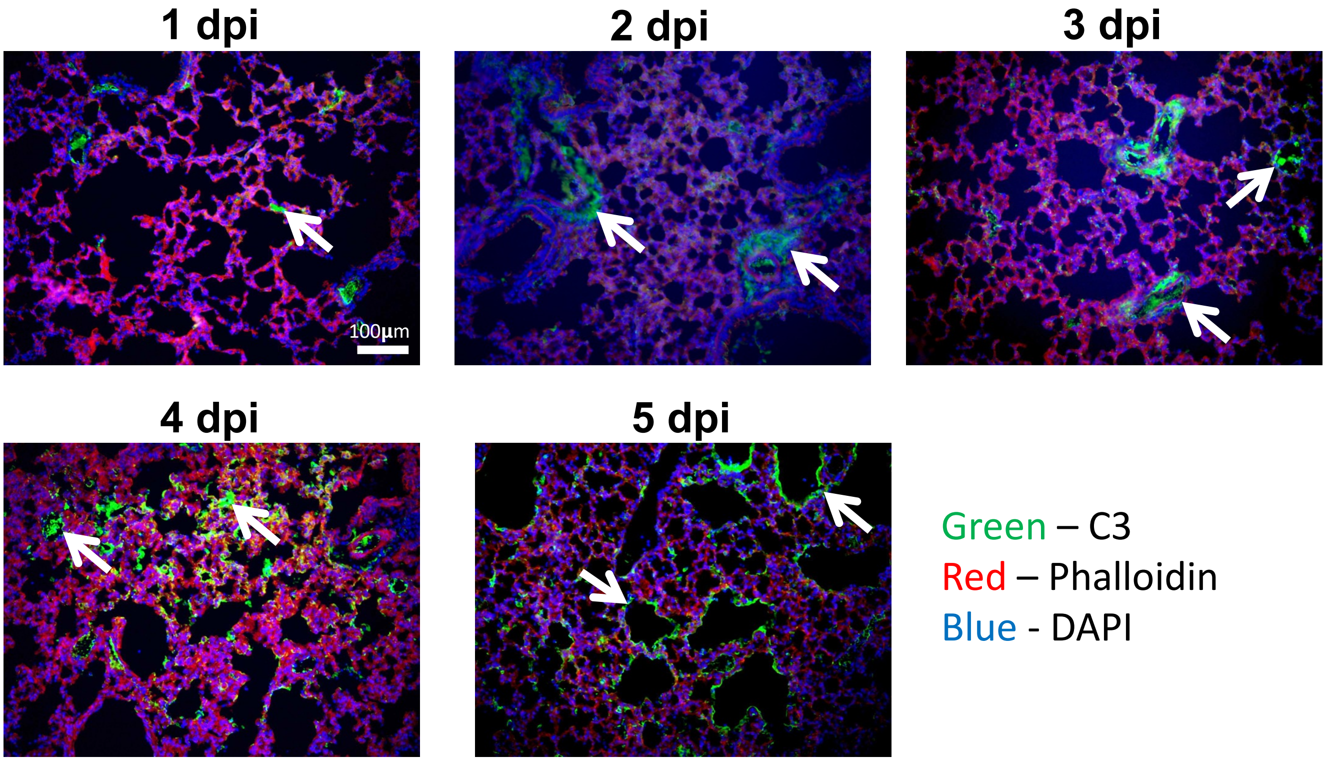


**Supplementary figure 3.** Complement C3 deposition in the lungs occurs early in SARS2-N501Y$_{MA30}$ infection. Serial immunofluorescence staining for C3 in BALB/c male mice following infection with SARS2-N501Y$_{MA30}$ demonstrates increased C3 deposition starting by 1 dpi, which progressively becomes more diffuse with time. White arrows indicate areas of C3 deposition within the lung. C3 = green, phalloidin = red, DAPI = blue.

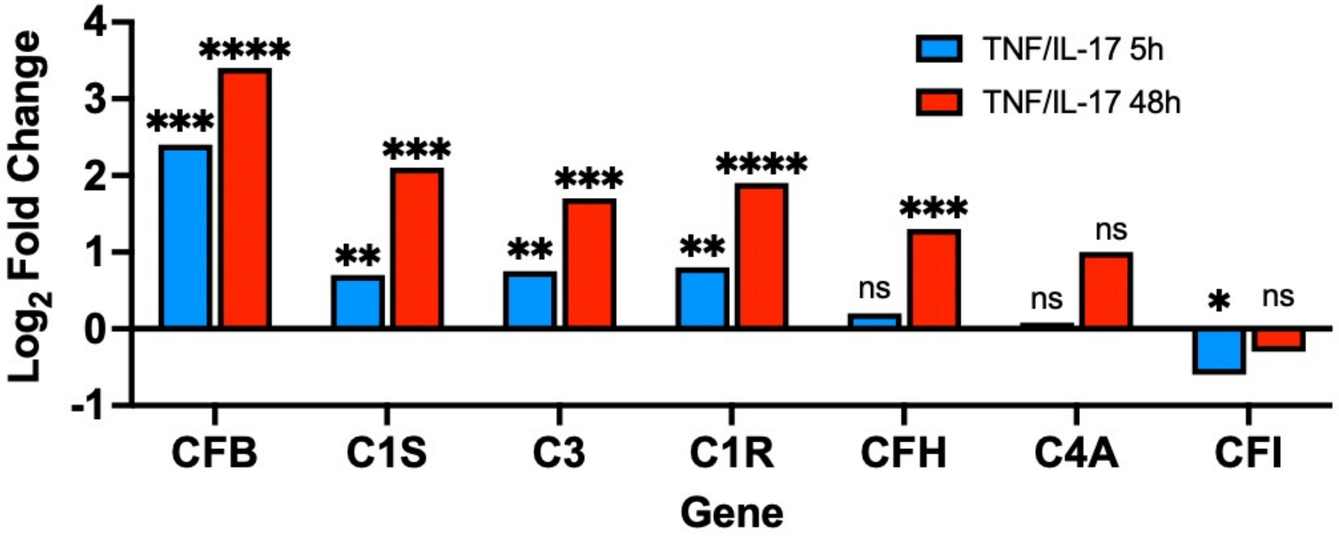


**Supplementary figure 4**. Complement gene expression increases in response to pro-inflammatory cytokines. Primary air liquid interface cultures of human airway epithelia (HAE) we treated with either vehicle or basolateral cocktail of IL-17 (20ng/mL) and TNF-$\alpha$ (10ng/mL) daily for 5 hours and 48 hours (n=6, normal donors). Results shown from bulk RNA sequencing demonstrate respiratory epithelial cells have an increased expression of genes associated with the complement cascade in the presence of inflammation. Each bar indicates the average $\log_2$ fold change for each gene with given treatment at the indicated timepoints. Significance determined by unpaired t-test. Where * p<0.05, ** p<0.001, *** p<1e-10, **** p<1e-20.

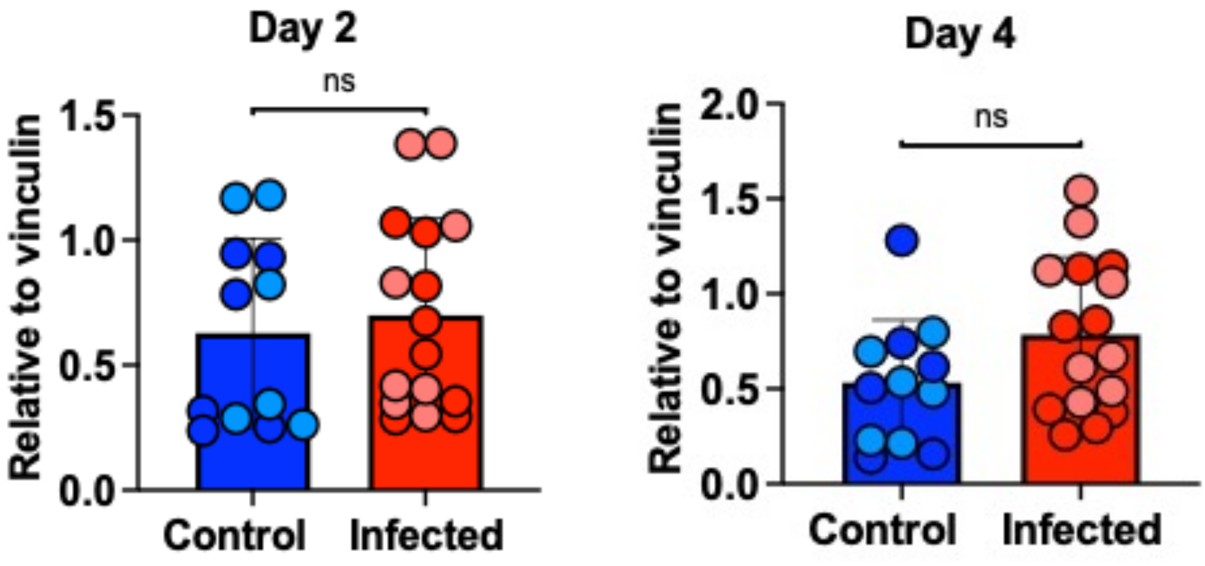


**Supplementary figure 5.** No evidence of C3 increase in kidney post-SARS2-N501Y$_{MA30}$ infection. Western blot densitometry for complement C3b on kidney homogenate from control (blue) and infected (red) on days 2 and 4 post-infection (*A* and *B*). Female mice indicated by light blue and light red dots, male mice indicated by cark blue and dark red dots. Significance determine by unpaired t-test with $p>0.05$. (n = 10).

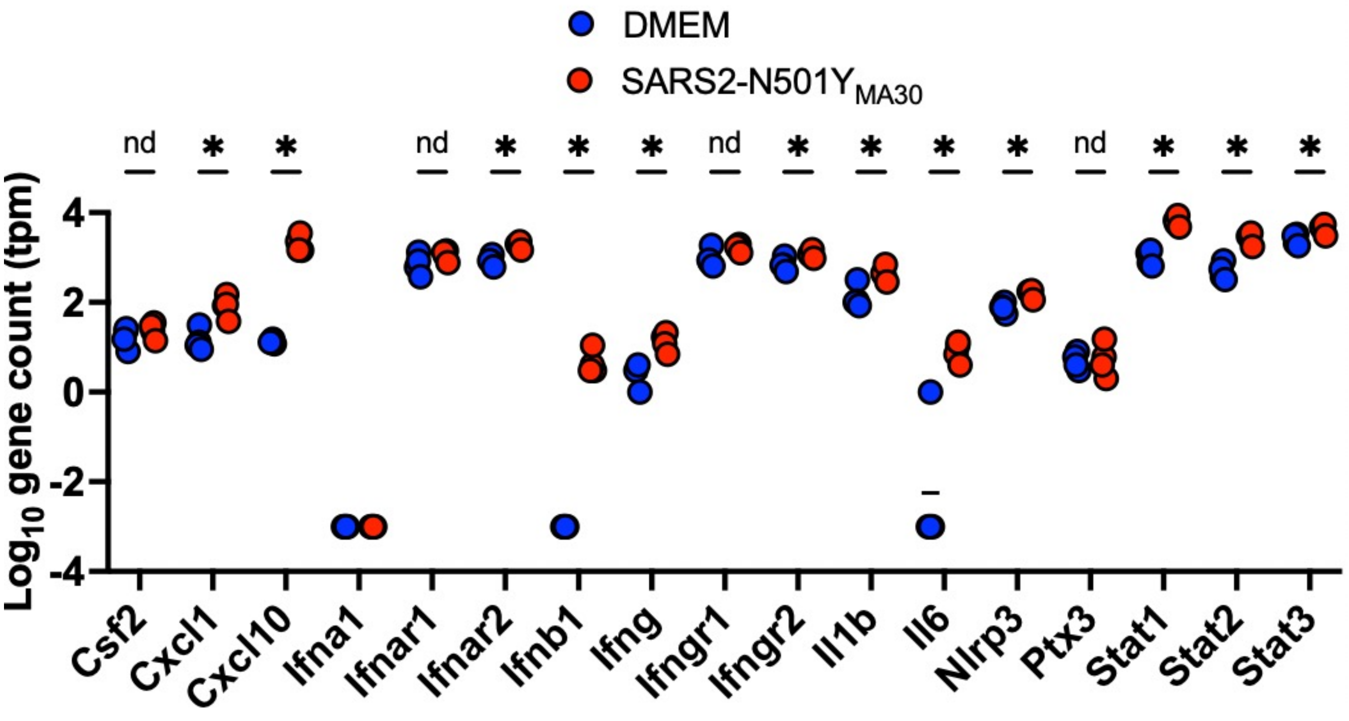


**Supplementary figure 6.** Log$_{10}$ gene counts as determined by bulk RNA sequencing for cytokine and chemokine expression in BALB/c mouse lung tissue on 5 dpi with 1,000 PFU of SARS2-N501Y$_{MA30}$ infection. Genes shown were selected from a list of those associated with either interferon mediated or complement mediated inflammation. Blue = DMEM, Red = SARS2-N501Y$_{MA30}$. n = 4. Significance determined by multiple unpaired t tests with * FDR<0.05, nd = no difference. Note, statistical significance unable to be determined for Ifna1 due to lack of expression.
